## Supporting information for "Emergence and maintenance of modularity in neural networks with Hebbian and anti-Hebbian inhibitory STDP"

### S1 Appendix. Untrained group of neurons.

This alternative protocol is analogous to the one of the numerical experiment reported in Fig. 1D of the main text. The only difference lies in the fact that a group of excitatory neurons is never stimulated and so it is untrained. The results obtained are described in Fig. S1. On the one hand, we obtain analogous results with the formation of two modular structures in the weighted connectivity associated to spontaneous recalls of the two different memories during the dynamical evolution as shown in raster plot. On the other hand, neurons of the untrained group are weakly connected among them in accordance with the absence of stimulation and are decoupled from the other clusters while receiving anti-Hebbian inhibition from them. As a result, these neurons spike in a totally asynchronous and irregular way, without impacting the dynamics of the rest of the network.

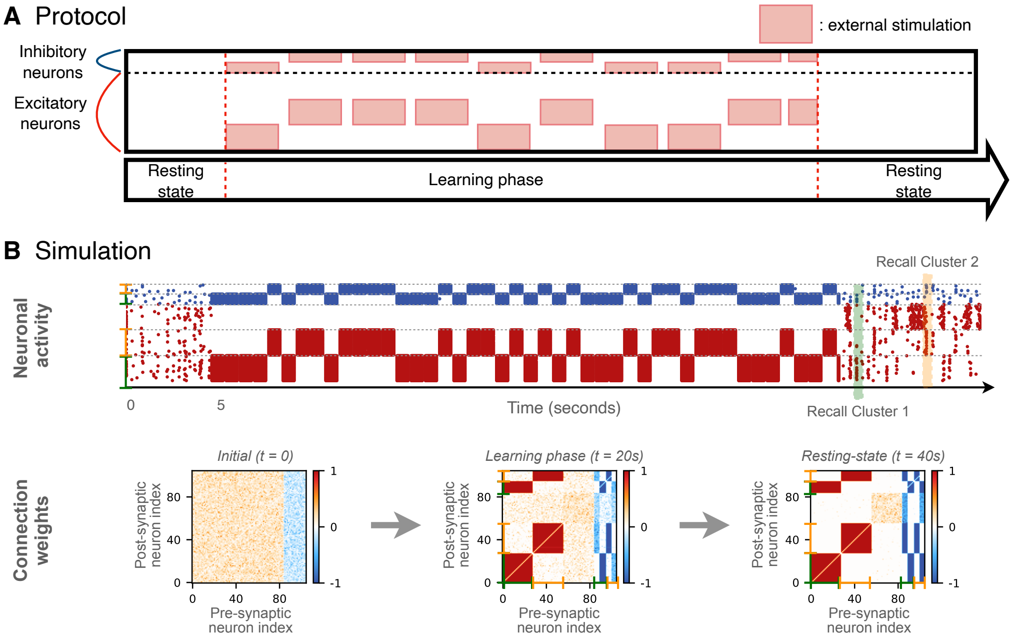


**Fig S1. Learning of 2 stimuli with an untrained sub-population. (A)** Stimulation protocol for a network of $N = 105$ neurons entrained with $M = 2$ stimuli with an untrained group of excitatory neurons. **(B)** Simulation and learning results. Connectivity matrices show the evolution of the synaptic weights leading to the emergence of two modules and the decoupling of the unstimulated sub-population. The raster plot shows the simulation for the three stages: initial resting phase, entrainment stage and the post-learning neuronal activity characterized by spontaneous recall events of $P_1$ neurons (green shadow) and $P_2$ neurons (orange shadow).

### S2 Appendix. Randomly stimulated neurons within each population.

This alternative protocol is analogous to that of the numerical experiment shown in Fig. 1D of the main text. The only difference lies in the fact that when a population is selected during learning, a random number of neurons in the excitatory population (with a probability of 0.5) is stimulated. The results obtained are described in Fig. S2. The direct consequence is that the two modular structures in the weighted connectivity are less well formed compared to the original experiment. Nevertheless, the clusters remain decoupled with the same feedforward and feedback inhibition described in the main text. Only the weights within the clusters appear have more random values. Therefore, although the spontaneous recalls of the two memory items are present in the dynamics as shown in the raster plot, they appears to be somewhat sparser and less synchronized. We assume that this is largely due to the fact that the connections to and from the inhibitory neurons are incomplete. In Fig. 3A of the main text, the randomness of the E-E connections does not impact the spontaneous recalls. This highlights the need to reach a convergence of the weights linked to inhibition to correctly memorize the items. Nevertheless, this experiment also shows that even if memory items are partially learned during each period of stimulation, the entire original memory item is somehow retrieved and learned.

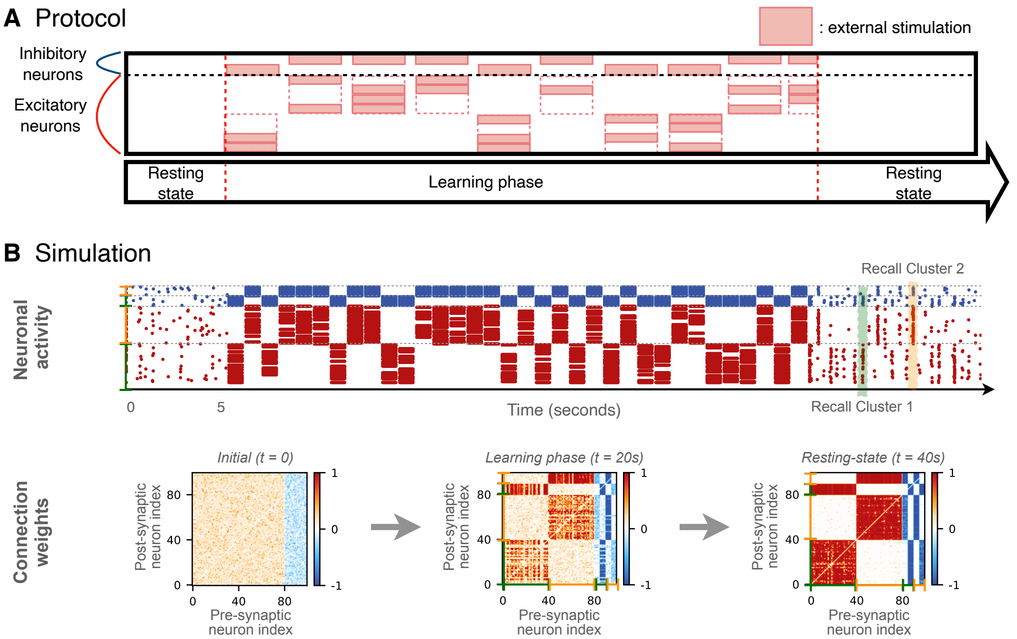


**Fig S2. Learning of 2 stimuli with randomly stimulated neurons within each distinct population.** **(A)** Stimulation protocol for a network of $N = 100$ neurons entrained with $M = 2$ stimuli, neurons within each population being randomly selected. **(B)** Simulation and learning results. Connectivity matrices show the evolution of the synaptic weights leading to the emergence of two modules. The raster plot shows the simulation for the three stages: initial resting phase, entrainment stage and the post-learning neuronal activity characterized by spontaneous recall events of $P_1$ neurons (green shadow) and $P_2$ neurons (orange shadow).

### S3 Appendix. Random stimulation values.

This alternative protocol is again analogous to that of experiment of Fig. 1D of the main text. The only difference lies in the fact that when a population is stimulated (excitatory and inhibitory neurons), the neurons within it receive inputs of random amplitude (i.e. inducing firing activity between 50 and 100 Hz). The results obtained are described in Fig. S3. We observe very similar results to those in Fig. S2. However in this case, the weight connections seem stronger than in the previous experiment. As a result, the spontaneous recalls appear to be more visible in the dynamics of raster plot. In addition to the conclusions made in the previous study, this experience allows us to conclude that the spatio-temporal correlations of the applied inputs are more impactful on the formation of the modular structures than the intensities of these same inputs. Indeed, the firing frequency induced by the inputs applied to the neurons necessarily has an impact on the encoding of information and the construction of memory as can be seen here. Nevertheless, compared with the previous experiment where the neurons received the same inputs but at times that were not necessarily correlated, the structure is much better learned in the current case.

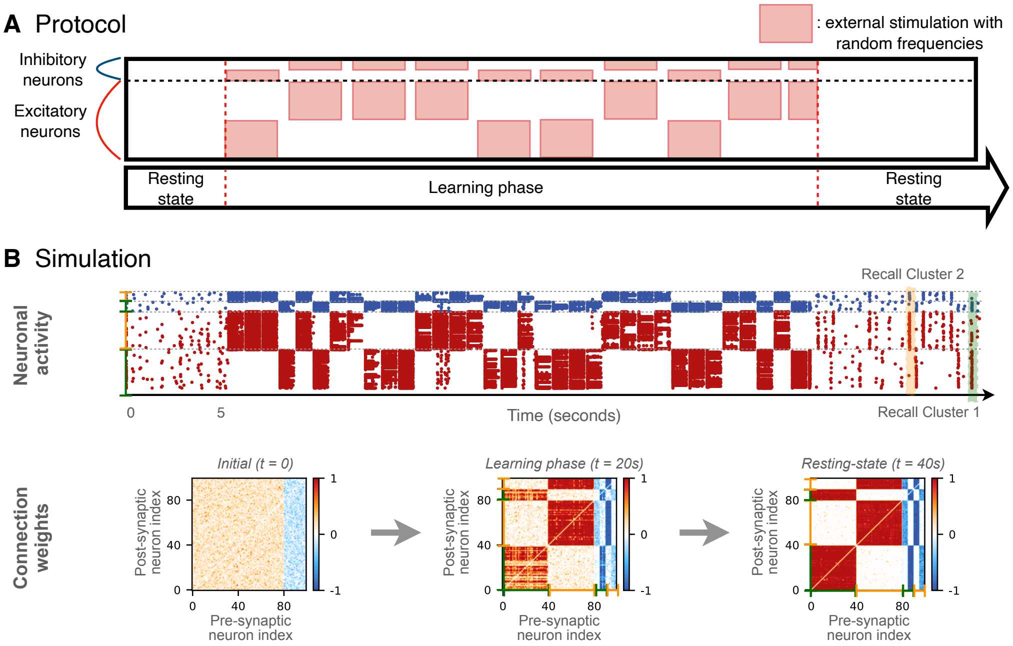


**Fig S3. Learning of 2 stimuli with random frequencies. (A)** Stimulation protocol for a network of $N = 100$ neurons entrained with $M = 2$ stimuli of random amplitude. **(B)** Simulation and learning results. Connectivity matrices show the evolution of the synaptic weights leading to the emergence of two modules. The raster plot shows the simulation for the three stages: initial resting phase, entrainment stage and the post-learning neuronal activity characterized by spontaneous recall events of $P_1$ neurons (green shadow) and $P_2$ neurons (orange shadow).

### S4 Appendix. Stability of four structural modules.

The reported numerical experiments analyse the limiting cases concerning the stability of four structural modules in absence of any stimulation. In Fig. S4A, we consider the case where each population contains only one Hebbian and one anti-Hebbian inhibitory neuron (i.e. a total of $N_I = 2 \times 4 = 8$ inhibitory neurons). This arrangement corresponds to the upper limit for the number of inhibitory neurons needed to maintain 4 independent memory items, represented by the red line in Fig. 4B of the main text. We observe that these conditions are sufficient for each cluster to present distinct spontaneous recall in neuronal activity.

In Fig. S4B, we break this limit by allocating only an Hebbian inhibitory neuron to the population $P_1$. We find that even if other populations are correctly recalled, memory recall of cluster 1 can occur at similar instant to that of the other clusters. Indeed, during recall, population $P_1$ does not inhibit the activity of the other populations, letting them activate. This has a direct effect on the consolidation process, where simultaneous recall of two memories patterns tends to induce their structural merging.

In Fig. S4C, we perform an opposite test, assigning only an anti-Hebbian inhibitory neuron to the population $P_1$. In that case, we observe a very short period of spontaneous activity with recalls from the other population. However, once population $P_1$ becomes active, it totally dominates the others, inhibiting them. These results are very similar to those observed in Fig. 1B. Indeed the absence of feedback inhibition provided by Hebbian inhibitory neuron, prevents $P_1$ activity to be regulated. Ultimately, this has no direct impact on the long-term maintenance of memory items in the weight matrix since their activity is suppressed. Nevertheless, this causes issues in the processing of stored information, since the network is blocked in an abnormal state.

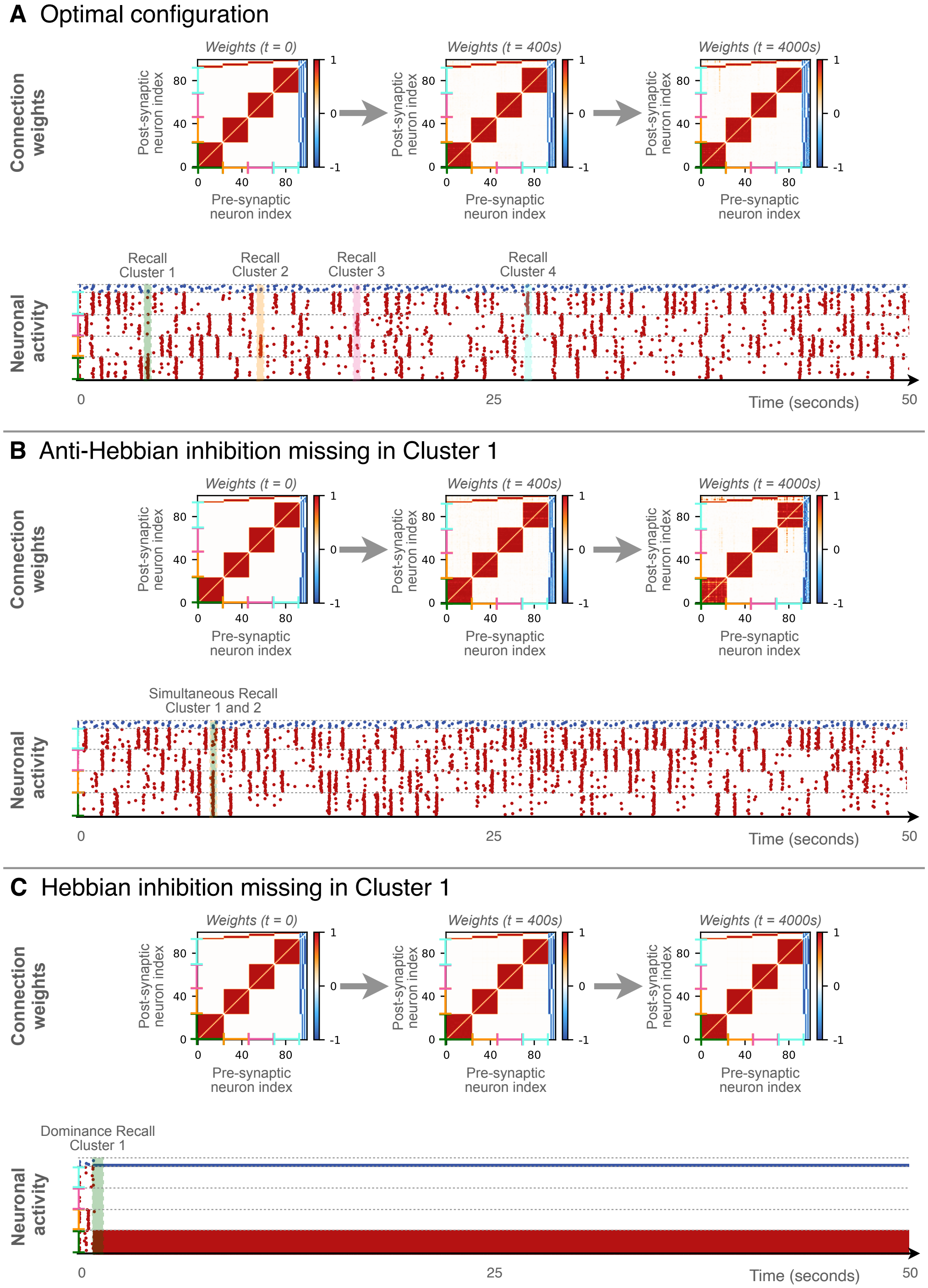


**Fig S4. Evolution of a network initially made of four structural modules in absence of any stimulation.** **(A)** Network of 92 excitatory neurons and 8 inhibitory neurons (one Hebbian and one anti-Hebbian in each cluster). **(B)** Network of 93 excitatory neurons and 7 inhibitory neurons (only an Hebbian in cluster 1). **(C)** Network of 93 excitatory neurons and 7 inhibitory neurons (only an anti-Hebbian in cluster 1). In each case, we study the stability of the organization from a structural and dynamical point of view. In each panel, the connectivity matrices show the evolution of the synaptic weights and the raster plot shows the neuronal activity. The green, orange, pink and cyan brackets and shadows represent clusters 1, 2, 3 and 4.

### S5 Appendix. Four overlapping stimuli.

In this alternative protocol, we reproduce the experiment of Fig. 6 of the main text but considering $M = 4$ stimuli which share 8 neurons. The results obtained are described in Fig. S5. We obtain similar results as in main text with the formation of four modules in the connectivity matrix as expected, together with the formation of hubs which are here connected (incoming and outgoing connections) with the four clusters. Regarding the dynamics in raster plot, the resting-state activity is comparable that observed in Fig. 6. We identify different types of spontaneous recalls involving one of the four clusters alone, recalls of one cluster accompanied by the hubs, or recall events of hubs alone. This experiment again highlights the richness of the different dynamics that the network can display and maintain for a certain period of time.

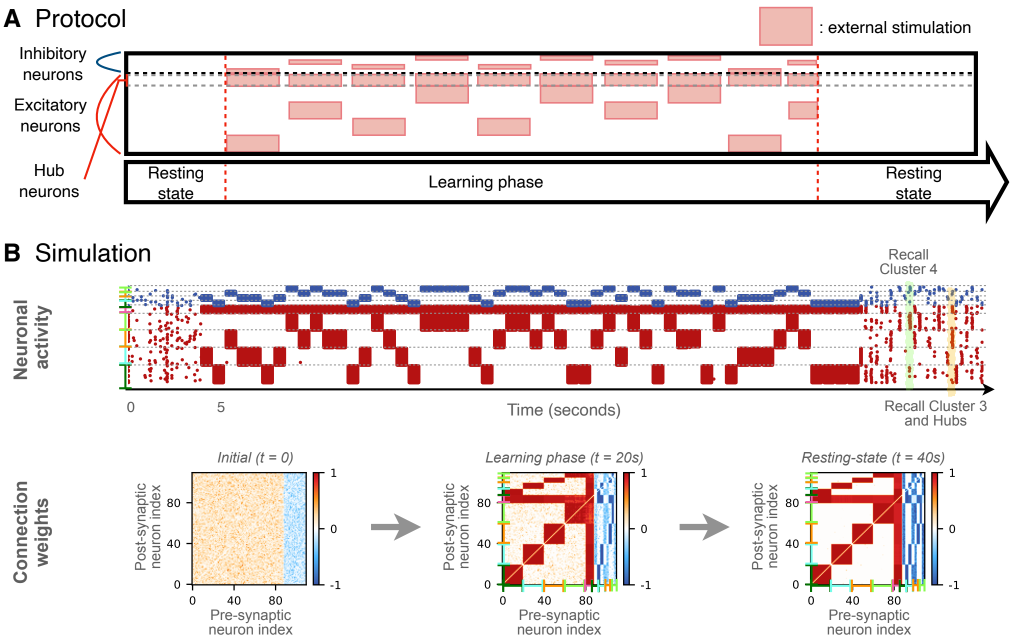


**Fig S5. Learning of 4 overlapping stimuli. (A)** Stimulation protocol for a network of $N = 110$ neurons entrained with $M = 4$ stimuli that share 8 excitatory neurons. **(B)** Simulation and learning results. Connectivity matrices show the evolution of the synaptic weights leading to the emergence of four modules which overlap over 8 hub neurons. The raster plot shows the simulation for the three stages: initial resting phase, entrainment stage and the post-learning neuronal activity characterized by a variety of spontaneous recall events as of $P_3$ neurons with hubs (orange shadow) or $P_4$ neurons without hubs (green shadow).

### S6 Appendix. Large and sparse networks.

In order to validate the stability of the model for larger network size and sparser connectivity, we apply the protocol reported in Fig. 1D of the main text to networks of larger size and with randomly connected neurons, see Fig. S6A.

Firstly, we consider a network that is initially globally coupled but with random synaptic weights and additive Gaussian noise, of $N = 20000$ neurons, where we still have 80% excitatory and 20% inhibitory neurons. The results of the experiment are reported in Fig. S6B. Secondly, we consider a random Erdös-Reniy network composed of $N = 1000$ neurons (where we still have a ratio $4 : 1$ for excitatory versus inhibitory neurons), with a probability of 50% of possible directed connections between neurons. In this case, we omit the additive Gaussian noise to check whether the random and sparse connectivity introduces sufficient heterogeneity in the synaptic inputs of the neurons to induce an asynchronous irregular behaviour. The results of this further experiment are reported in Fig. S6C.

Qualitatively, we obtain the same results as in Fig. 1D, with the formation of two modular structures in the weight matrix joined to spontaneous recalls of the two different memories during the post-learning phase. The main differences are indeed observable in this regime. In Fig. S6B, due to the size of the network it is evident that a large number of neurons fires randomly and independently, this renders more difficult to identify the spontaneous recall events. However, they still take place and the memory should be consolidated on the long term as shown in Fig. 5 of the main text.

In the case of the sparse network, the activity during the resting state also presents an irregular activity of the neurons despite the dynamical behaviour is deterministic. However, also in this case, as shown in Fig. S6C, the modular structures emerge in the weight matrix. One should notice that the orange color in the weight matrix at $t = 50$ seconds is due to the fact that 50% of the neurons are disconnected and do not reflect to the actual value of the weights, that have the same values as in Fig. S6C.

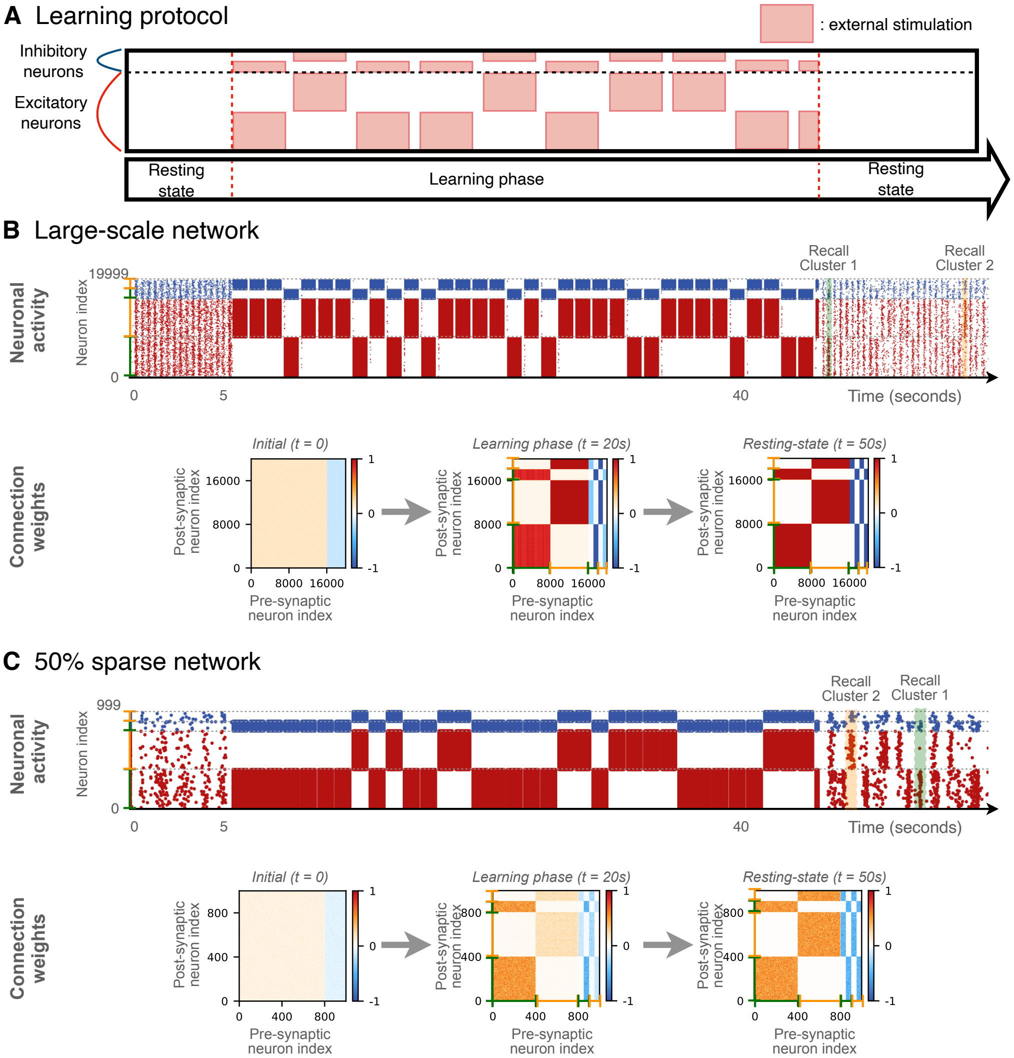


**Fig S6. Model validation in larger and sparser networks. (A)** Stimulation protocol for a network entrained with $M = 2$ stimuli. **(B)** Simulation and learning results for $N = 20000$ neurons. Connectivity matrices show the evolution of the synaptic weights leading to the emergence of two modules. The raster plot shows the simulation for the three stages: initial resting phase, entrainment stage and the post-learning neuronal activity characterized by spontaneous recall events of $P_1$ neurons (green shadow) and $P_2$ neurons (orange shadow). **(C)** Simulation and learning results for $N = 1000$ neurons and 50% sparse connection without additive noise terms. Connectivity matrices show the evolution of the synaptic weights leading to the emergence of two modules. The lighter color is due to the sparsity of the connections, but the numerical values are the same as in panel (B). The raster plot shows the simulation for the three stages: initial resting phase, entrainment stage and the post-learning neuronal activity characterized by spontaneous recall events of $P_1$ neurons (green shadow) and $P_2$ neurons (orange shadow).

### S7 Appendix. Storing the maximum number of items.

Here, we reproduce the experiment of Fig. 4A of the main text, but considering $M = 33$ stimuli. This limiting case corresponds to the maximum capacity of items that can be learned and recalled for a network of $N = 100$ neurons (marked by the star symbol in Fig. 4B). This implies considering a network composed of $N_I = 66$ inhibitory neurons (with 33 anti-Hebbian and 33 Hebbian). Except these differences in the initial conditions, the network is trained as usual.

This protocol leads to the formation of 33 memories, where each item is composed of an excitatory neuron, a Hebbian and an anti-Hebbian inhibitory neuron, as shown in the connectivity matrix and in the diagram of Fig. S7B. Consequently, each anti-Hebbian inhibitory neuron projects to all the other clusters of neurons. Concerning the post-learning activity, given the small size of the clusters now, it is difficult to distinguish individual memory recalls with clarity. Nevertheless, the asynchronous dynamics of the network at rest tends to confirm that all excitatory neurons have a decorrelated activity. To visualize this, we estimated the instantaneous Kuramoto order parameter $R$ throughout the simulation. During the post-training resting state the system is essentially desynchronized, since $R \simeq 0.1 \simeq 1/\sqrt{N}$, as expected in an asynchronous system made of $N = 100$ neurons/oscillators due to the central limit theorem. On the contrary, before the training phase the system was partially synchronized, since $R > 0.6$. We have also defined a normalized spike count associated to a certain memory item

$$\rho(t) = \frac{n_m^{sp}(t)}{n^{sp}(t)}$$

where the spike count associated to the neurons related to a given memory item $n_m^{sp}(t)$ is divided by the total number of spikes emitted in the network $n^{sp}(t)$. By computing $\rho(t)$ for the memory highlighted in purple in the figure, we can identify clear recalls when $\rho = 1$, meaning that the spikes only come from the neurons associated to the recalled memory.

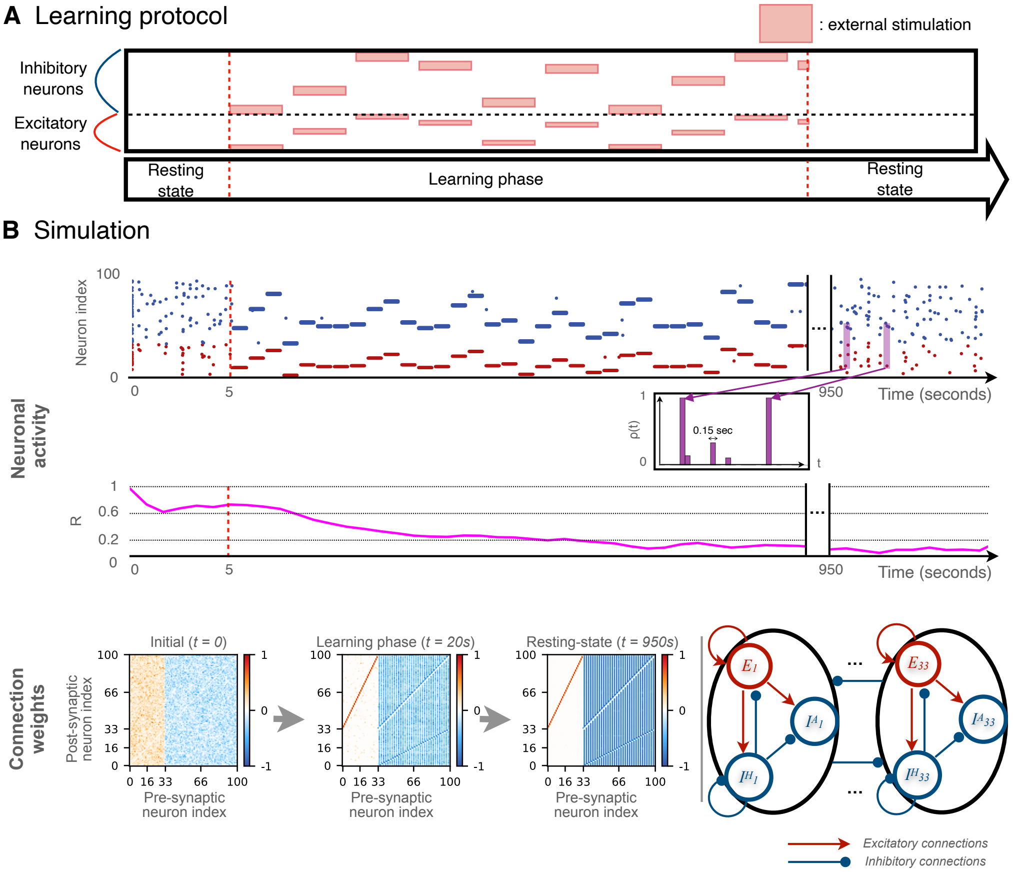


**Fig S7.** **Maximum network capacity.** **(A)** Stimulation protocol for a network of $N = 100$ neurons entrained with $M = 33$ stimuli. **(B)** Simulation and learning results. Connectivity matrices show the evolution of the synaptic weights leading to the emergence of 33 modules. The final configuration of the connection weights is shown schematically on the right. The raster plot shows the simulation for the three stages: initial resting phase, entrainment stage and the post-learning neuronal activity characterized by a variety of spontaneous recall events of the memories. The recalls of one memory are highlighted in purple with its normalised spike count $\rho(t)$ for bins of 0.15 sec. The instantaneous Kuramoto order parameters $R$ of the network throughout the simulation time is displayed bellow.

#### S8 Appendix. Stability of the stored items for increasing network sizes.

By following [1], we have analysed the stability of the intra- and inter-cluster weights for increasing system sizes and numbers of memory items (essentially restricted to $N = 100$ neurons and $M = 2$ memories in the main text). We have here adopted two protocols. In the first, the number of memory items is fixed to $M = 10$. In the other, $M$ grows proportionally with $N$, in this way the number of neurons associated to each memory clusters remains the same. In both cases we considered three different system sizes, namely, $N = 200$, 500 and 1000 neurons. The results obtained for $M = 10$ and by varying $N$ are shown in Fig. S8, while those corresponding to $M = N/100$ are reported in Fig. S9.

As a general result, we observe that the intra-cluster weights essentially remain constant over time independently of $N$ for both protocols, indicating that the internal stability of the formed clusters is independent of the system size. On the other hand, for the protocol shown in Fig. S8, we observe a tendency for the inter-clusters weights to grow/decrease over time during the post-learning phase, ultimately leading to a merging of the stored memory items. This can be explained by the fact that reducing network size, reduces the number of inhibitory neurons allocated to each memory, decreasing their stability. Indeed, even if the configurations are sufficient to guarantee coherent reactivations (see Fig. S7), less inhibition increases the probability of incoherent spikes due to network variability. However, the rate at which the inter-cluster weights modify strongly decreases with $N$, indicating that for larger system sizes the memories remains stable for longer times.

Finally, by considering the situation where the ratio $M/N$ is maintained constant as in [1], all intra- and inter-cluster weights remain essentially constant over 4 hours of post-learning evolution, displaying no drift in their values. By employing this second protocol we can extrapolate the memory capacity characterized in the main text for $N = 100$ in Fig. 4B to networks of larger size.

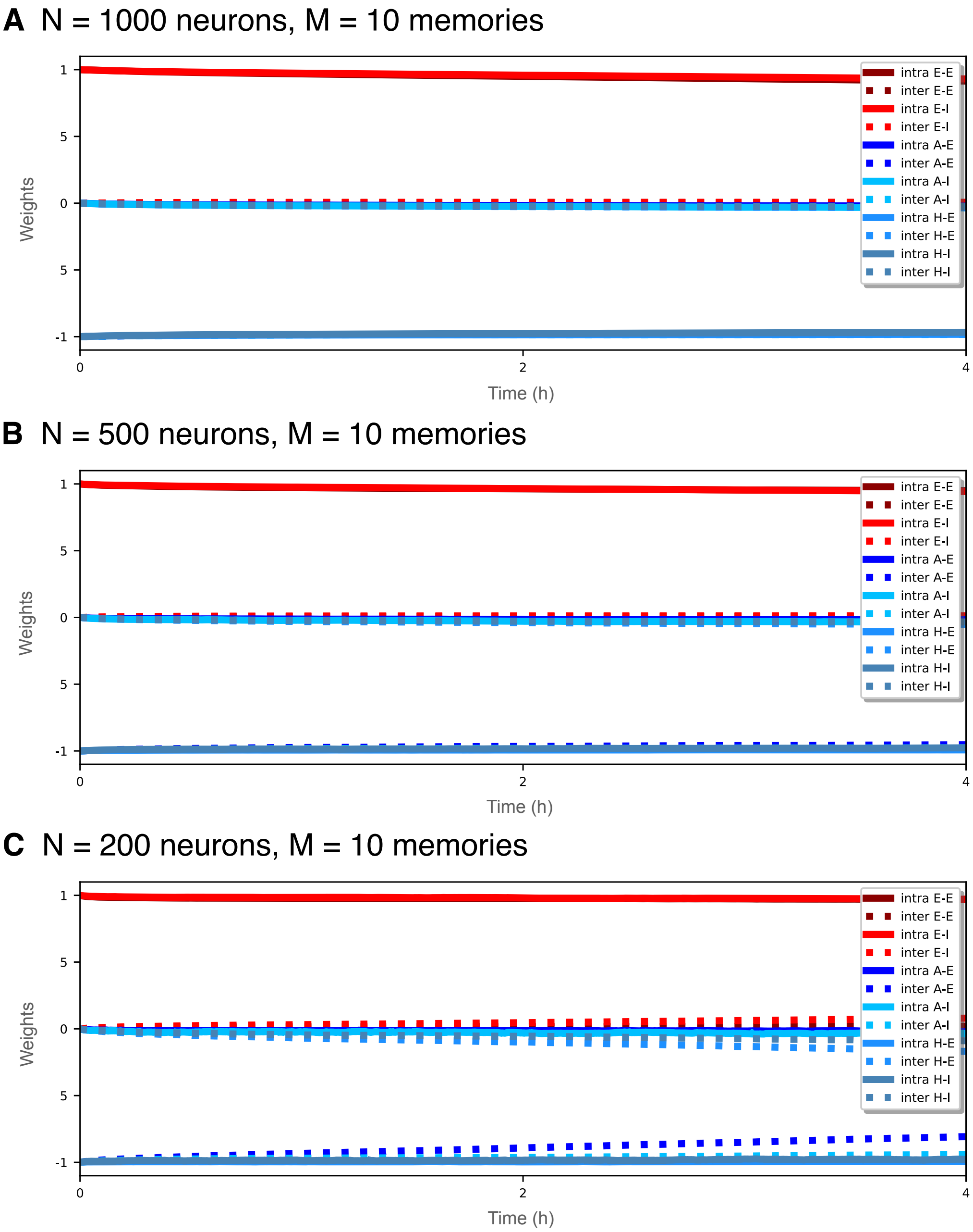


**Fig S8. Stability of the network connections scaling up the network size.** Post-learning evolution of mean intra- (solid lines) and inter- (dashed line) clusters weights for excitatory to excitatory (E-E dark red), excitatory to inhibitory (E-I red), anti-Hebbian inhibitory to excitatory (A-E dark blue), anti-Hebbian inhibitory to inhibitory (A-I cyan), Hebbian inhibitory to excitatory (H-E blue) and Hebbian inhibitory to inhibitory (H-I steel blue) connections; for **(A)** $N = 1000$ neurons, **(B)** $N = 500$ neurons, and **(C)** $N = 200$ neurons, all trained with $M = 10$ stimuli.

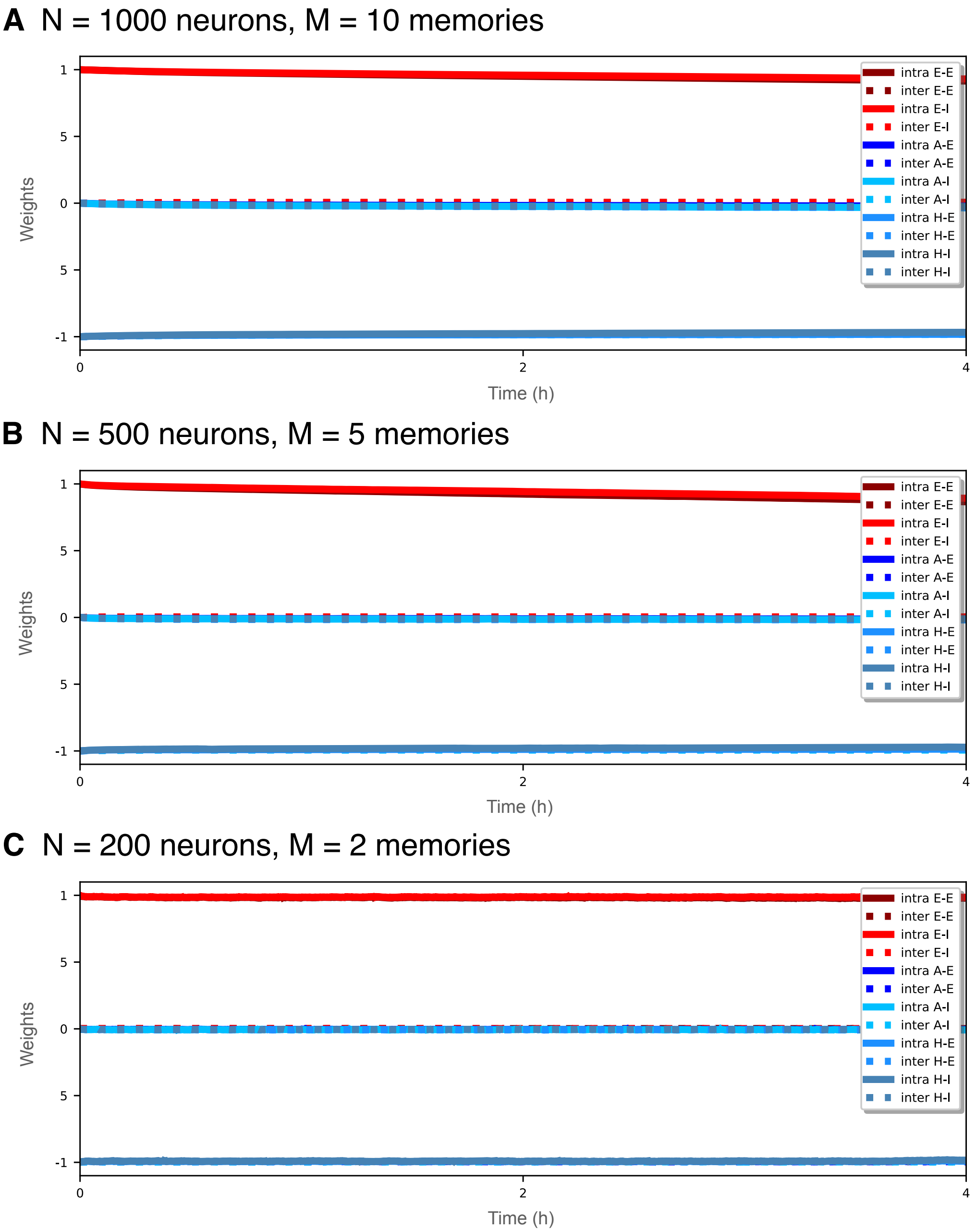


**Fig S9. Network connections stability scaling up $M$ proportionally to $N$.** Post-learning evolution of mean intra- (solid lines) and inter- (dashed line) clusters weights for excitatory to excitatory (E-E dark red), excitatory to inhibitory (E-I red), anti-Hebbian inhibitory to excitatory (A-E dark blue), anti-Hebbian inhibitory to inhibitory (A-I cyan), Hebbian inhibitory to excitatory (H-E blue) and Hebbian inhibitory to inhibitory (H-I steel blue) connections; for **(A)** $N = 1000$ neurons, **(B)** $N = 500$ neurons, and **(C)** $N = 200$ neurons, trained respectively with $M = 10$, $M = 5$, $M = 2$ stimuli.

### S9 Appendix. Estimation of the time needed to forget a recall event.

The rate of change of the synaptic weights depends on several factors: the current value of the weight $w_{ij}(t)$, the number of neurons spiking at a given time $t$, and the temporal precision of their spikes. Therefore, it is non-trivial to quantify the precise amplitude of the change of weights at all times. If a synapse is already at its maximum weight capacity, a potentiation (due to a recall) has almost no impact whereas the absence of recalls leads to more impactful depression (see soft bound functions in Methods, Figs. 7D and E).

We can derive an approximation for the impact of a recall on a synaptic weight—compared to a forgetting epoch—by inspecting closer the plasticity function (see Fig. 7A). For uncorrelated spikes during an asynchronous irregular firing epoch (i.e. for $|\Delta t| > 0.5$), the weights depress proportionally to $\Lambda(\Delta t) \approx -0.1$ due to the forgetting term. On the other hand, if the spikes are fully synchronized (i.e. $\Delta t = 0$), their increase is proportional to $\Lambda(\Delta t) = A_+ - A_- - f = 2.347 - f$. Thus, by considering an average weight of $w_{ij} = 0.5$ (where $\tanh(\lambda(1 - w_{ij})) = \tanh(\lambda w_{ij}) \approx 1$) Eq. 10 becomes:

$$\frac{[w_{ij}(t^+) - w_{ij}(t^-)]}{\gamma_l} \simeq \begin{cases} 2.347 - f, & \text{for synchronized spikes during a recall} \ , \\ -f, & \text{for uncorrelated spikes during AI state} \ . \end{cases} \quad \text{(S1)}$$

The ratio of these two (absolute) values for potentiation and depression for $M$ stimuli and $f = f_0/M$ is: $\frac{2.347}{f_0}M - 1 = 11.735M - 1$, where $f_0 = 0.2$, and it gives an estimate of the number of uncorrelated spikes that would be required to lose the acquired potentiation. The number of these spikes grows linearly with $M$. By considering an average firing rate of 2 Hz for the uncorrelated firings, this implies that a period of $\simeq 11$ seconds would be required to forget the contribution of a single recall event for $M = 2$, that will grow to $\simeq 58$ seconds for $M = 10$. The prolongation of this time window proportionally to $M$ will allow to all the stored memory items to be randomly recalled before they are forgotten.
